# Long-Range Glutamatergic Projections Target Hippocampal Cajal-Retzius Cells Through Atypical Receptor Signaling

**DOI:** 10.64898/2026.10.16.759635

**Authors:** Kristian Moan, Giulia Quattrocolo

## Abstract

Cajal-Retzius (CR) cells are a population of early-born, transient neurons that serve an indispensable role during embryonic brain development. Postnatally, most CR cells undergo programmed cell-death, except in a few areas, including the hippocampus, where they persist until adulthood. The hippocampal population of CR (hCR) cells have previously been implicated in receiving and facilitating ingrowth of entorhinal afferents and the establishment of the hippocampal circuit. Nevertheless, only local GABAergic synaptic input has been demonstrated onto hCR cells before. Through transsynaptic rabies tracing we show that hCR cells receive input for at least seven extrahippocampal areas, all putative glutamatergic populations, even though CR cell spontaneous activity is speculated to be exclusively mediated through GABAergic action. Here, we show that a portion of CR cell spontaneous activity is strychnine-sensitive, and that glutamatergic fibers from entorhinal cortex layer II apposition themselves in physiological distance (<300 nm) to glycinergic receptors on hCR cells. We propose that hCR cells guide several, if not all, afferents terminating in stratum lacunosum moleculare and the outer molecular layer in the hippocampus. These findings shine new light on how hCR cells might facilitate hippocampal maturation.

## Introduction

Cajal-Retzius (CR) cells are early generated, transient neurons located throughout neocortical and allocortical areas, specifically in layer (L) 1, and the molecular layer (ML) of dentate gyrus (DG), and stratum lacunosum-moleculare (SLM) of the Cornu Ammonis (CA)-areas in hippocampus (Causeret et al., 2021; de Frutos et al., 2016; Elorriaga et al., 2023; Moan et al., 2025; Soriano et al., 1994). CR cells hold an essential role in early brain development as they facilitate correct layer alignment of migrating neurons (Ogawa et al., 1995) and layer specific termination of entorhinal fibers to the hippocampus (Ceranik et al., 2000; Del Río et al., 1997). In later postnatal periods, they refine local circuits in the neocortex (Damilou et al., 2024; Riva et al., 2023) and the hippocampus (Glærum et al., 2024). Specifically in the hippocampus, early ablation of CR cells impedes an animal’s ability to learn (Anstötz et al., 2022; Glærum, 2024) and disrupts processing of spatial navigation signals (Sachuriga et al., 2026). In the murine brain, CR cells disappear from most of the neocortex shortly after birth, with 80 - 97% gone by the second postnatal week (Chowdhury et al., 2010; del Río et al., 1995; Moan et al., 2025). They persist into postnatal development and even adult life in the hippocampus and parts of the parahippocampal region (Anstötz et al., 2018; Del Rıo et al., 1996; Moan et al., 2025; Supèr et al., 1998), with up to 20 - 30% retention in the hippocampus at postnatal day (P) 15. The prolonged survival of hippocampal Cajal-Retzius (hCR) cells suggests an ongoing maturation of hippocampal networks. Accordingly, insight into their embedding in hippocampal circuitry will help us elucidate hippocampal network functions, both in normal and dysfunctional development.

Postnatal hCR cells take part in the local circuit of hippocampal subfields CA and DG. They are glutamatergic neurons which generate output onto pyramidal neurons and neurogliaform (NGF) cells, in CA1. Locally, their input originates from NGF cells and Oriens Lacunosum-Moleculare (OLM) cells, creating what appears to be an exclusively GABAergic drive (Quattrocolo & Maccaferri, 2013, 2014). This is corroborated by the full abolishment of spontaneous current events after application of GABA receptor antagonists (Cosgrove & Maccaferri, 2012; Kilb & Luhmann, 2001; Marchionni et al., 2010; Soda et al., 2003). At the same time, other studies have demonstrated that CR cells are targeted by glutamatergic fibers (Anstötz et al., 2014), likely from outside of the hippocampus (Del Río et al., 1997; Supèr et al., 1998), a puzzling observation, as CR cells show almost no response to stimulation with glutamate (Marchionni et al., 2010). Notably, recorded spontaneous postsynaptic currents from CA1 CR cells show certain events with kinetics and amplitudes that do not conform to those mediated by NGF- or OLM- inputs (Quattrocolo & Maccaferri, 2013), but are likely GABAergic in nature. Therefore, to appreciate the full extent of the CR cell’s role in the hippocampal network, we sought to recognize all sources, and the potential nature, of their inputs.

Results from pseudotyped rabies (RABV) injections at P1 demonstrated that hCRs receive input from at least 7 extrahippocampal regions, all from putative glutamatergic populations. Super-resolution imaging revealed that glutamatergic fibers from L II medial entorhinal cortex (MEC) fibers position themselves in a physiologically plausible distance (<300 nm) from glycinergic receptors on hCRs, indicating that these fibers might communicate through delivery of glycine. This possibility was corroborated by in-vitro patch clamp experiments, where we observed the frequency of spontaneous postsynaptic current events increasing after taurine application, and frequency and amplitude decreasing after strychnine application.

## Material and Methods

### Animals

In total 21 animals were used for experiments in this project. All experiments were conducted in compliance with protocols approved by the Norwegian Food Safety Authorities and European Directive 2010/63/EU (FOTS ID 24847; FOTS ID 31016). Pups were separated from the mother at postnatal day 21. Animals were housed with up to five animals per cage and kept in an inverted 12-hour light/dark cycle, with enriched cages and food and water ad libitum. 12-24 hours before surgery, animals were separated into individual cages. If possible, they were housed back together with littermates after surgery.

For retrograde transsynaptic tracing experiments, Pde1c-cre transgenic mice (B6.FVB(Cg)-Tg(Pde1c-cre)IT146Gsat/Mmucd, MMRRC 030708) (Osheroff & Hatten, 2009) were bred with NDNF-flox-FlpO transgenic mice (donated by Dr. Rob Machold, NYU and Dr. Gord Fishell, Harvard University (Chamberland et al., 2024; Moan et al., 2025)), creating Pde1c-Cre+/−;NDNF-flox-FlpO+/− mice, used as the experimental group (*n* = 3) and their double-negative littermates were used as the control group (*n* = 3). For in-vitro electrophysiology experiments, Pde1c-cre^+/−^;NDNF-flox-FlpO^+/−^ animals were bred with homozygous Ai65 (B6.Cg-Gt(ROSA)26Sortm65.2(CAG-tdTomato)Hze/J) (JAX stock #021875) animals to produce Pde1c-Cre+/−;NDNF-flox-FlpO+/−;Ai65+/− (*n* = 14). For EC LII input to hCR cell tracing, we crossed Pde1c-cre transgenic mice with Odz3-tta (originally published as MEC-13-53A (Blankvoort et al., 2018)) and tetO-GCamp6 mice (Blankvoort et al., 2018), to create Pde1c-cre^+/−^;Odz3-tta^+/−^; tetO-GCamp6^+/−^ transgenic animals (*n* = 1). The respective mouse line enables access to CR cells with Cre-dependent vectors and GCamp6 expression in entorhinal LII reelin-positive cells. Even though the animals expressed GFP through the GCamp, we decided to inject the virus to achieve a stronger signal. Animals of both sexes were used for experiments.

### Surgery

A total of 7 animals were injected for viral labelling. Animals were unilaterally injected at either P1 or P30. For the retrograde transsynaptic pseudotyped rabies experiments, the vector (RabV CVS-N2c(DeltaG)-tdTomato) (RabV) was mixed at 1.66:1 ratio with the recombinant adeno-associated virus (AAV2/1-CAG-fDIO-TVA-GFP-G) (helper virus) and injected as a cocktail in P1 animals. We performed one injection at P30 for a MEC LII input tracing experiment, when 150 nl of AAV2/1-flex-mCh was injected in the hippocampus, while 2x 200 nl of AAV2/1-TetO-GFP was injected at two different dorsoventral coordinates in the MEC of a Pde1c-cre^+/−^;Odz3-tta^+/−^;tetO-GCamp6^+/−^ animal.

### Viral injections at P1

The animal was placed on a heating pad (37 °C) and mounted in a stereotaxic frame (David Kopf Instruments, Tujunga CA, USA) with a custom-made adaptor. A piece of tape with a diamond cutout was used to fix the head and stretch the skin. Anesthesia was induced with 5% isoflurane while maintained with 3% (airflow 1L/min, Baxter AS). Coordinates for injections were calculated based on *lambda* (AP: +0.8 mm; ML: ±1.2 mm; Z: −1.22 mm) for each animal. A glass capillary filled with the necessary amount of virus was prepared immediately before anesthesia. First, capillaries (Drummond Scientific Company) were made on a pipette puller (Flaming/Brown Micropipette puller, Model P-87, Stutter instruments co., USA). The tip of the capillary was cut off with small surgical scissors, before being filled with mineral oil (Merck Life Sciences AS/Sigma Aldrich Norway AS, Oslo, Norway). Subsequently, the amount of virus needed for the injection ∼200nl extra was pulled into the capillary. The glass capillary was placed on top of the head, at the right coordinates before it was lowered down into the brain. The injections were made with a Nanoject III injector (Drummond Scientific Company). For each animal 50 nl (only helper virus) or 100 nl (helper virus and RabV) were injected (stepwise injections, 10 nl volume, 0.10 nl/sec, with 2 seconds delay between steps). After finishing the injection, we waited 30 seconds to ensure complete diffusion of virus before retracting the capillary. The animal was placed to recover on a heating mat (37 °C), before being placed back into home cage.

### Viral injections at P30 - adult

Animals were anesthetized with 4% isoflurane (airflow 1L/min, Baxter AS) and transferred to a small animal stereotaxic frame with surgical mask (David Kopf Instruments, Tujunga CA, USA), and heating pad (37 °C). Following, meloxicam (1 mg/kg, SC, Metacam, Boheringer Ingelheim Vetmedica) and buprenorphine-hydrochloride (0.01 mg/kg, SC, Temgesic, Indivior) were administered subcutaneously (SC). Bupivacaine hydrochloride (1 mg/kg, Marcain, Astra Zeneca), a local anesthetic, was injected subcutaneously (SC) over the incision site. The dorsal surface of the head was shaved, spanning from between the eyes to between ears, with an electrical shaver. The skin was washed with sterile NaCl (9mg/ml, B. Braun) and disinfected with 70% ethanol (Rectified ethanol, KiiltoClean AS) and Chlorhexidine (5mg/ml, Fresenius Kabi). To expose the skull, a rostrocaudal incision was made. Next, to measure that the head was leveled, the dorsoventral level was measured over points bregma and lambda. Bregma was additionally used to measure the anterior-posterior coordinates, while the midsagittal sinus was used for the mediolateral measurements. The surface of the dura was used to calibrate the dorsoventral measurements. A surgical drill attached to the surgical frame, with a drill head of 0.5mm, was used to calculate and navigate to the site of injection (Dorsal Dentate Gyrus, bregma as AP 0 and midsagittal sinus as ML 0: AP: −2.5mm, ML: −1.5mm, DV: −1.9mm; MEC, transverse sinus as AP 0 and midsagittal sinus as ML 0: AP: +0.5 mm, ML: 3.1 mm, DV: −1.9 mm + 1.8 mm). Next, a small hole was drilled over the site of injection before replacing surgical drill with injector and viral capillaries. Capillaries were prepared as described above, only cutting lower down, ensuring a smaller tip. The capillary was positioned over the dura and lowered down into the brain. Before starting injection, we waited 1 min. Injection was delivered as described above, but for adult animals we waited 10 min after finishing the injection to ensure complete diffusion of virus. After retracting the capillary, the wound was sutured close and washed with sterile NaCl. The animals were given a subcutaneous injection of 0.25 ml NaCl for hydration before being placed back in home cage, placed in a recovery rack (37 °C) shielded from sound and vibration. Animals received painkillers following surgery for up to two days after, with buprenorphine-hydrochloride administered orally, mixed with chocolate spread, first 6-8h after surgery and later every 12h, and meloxicam SC every 24h. To control healthy recovery their weight was measured every day until it stabilized at pre-operative level, or they gained weight, measured for at least three days. Animals housed together, that were injected on the same day, were housed back together if possible.

### Perfusion and section preparation

14 days after being injected with RabV the animals were sacrificed. Animals were first anesthetized with 0.1 ml isoflurane in a closed chamber, and injected with a lethal dose of pentobarbital (100mg/kg, 0.1ml, IP, SANVIO Pharma AS). They were transcardially perfused, with ice-cold (4 °C) phosphate-buffered saline (PBS) placed on ice, for 5 min (3.5 ml / min), before switching to fresh, ice-cold (4 °C) 4% paraformaldehyde (PFA, pH 7.4, Merck Life Sciences AS), for 6 min (3.5 ml / min). The brains were then removed from the skull and post-fixated in PFA for 16-24h at 4 °C. After post-fixation the brains were stored in 15% sucrose and finally 30% sucrose as cryoprotection in preparation for microtome sectioning. Brains were cut on a freezing microtome (Sliding microtome HM 430, Thermo Scientific, Walldorf, Germany) at −33.5 °C with 50 μm sections, in three series. Tissue was either stored temporarily in PBS at 4 °C or in anti-freeze solution (40% 125 mM PBS, 30% glycerol, 30% ethylene glycol) at −20 °C for long-term storage.

### Immunohistochemistry

Thin sections were first washed in PBS 3 × 10 min before blocking in solution containing 10% normal donkey serum (Jackson ImmunoResearch,) and 0.1% Triton X-100 (Merk) dissolved in PBS, for 30 min. After blocking sections were incubated in primary antibodies, in incubation solution containing 1% normal donkey serum and 0.1% Triton X-100 dissolved in PBS, for three overnights at 4 °C while rotating. Primary antibodies: Chicken anti-GFP 1:1000 (Abcam, ab13970), Guinea pig anti-NeuN 1:1000 (Merk, ABN90P), Goat anti-mouse Reelin 1:500 (Biotechne RandD systems, AF3820), Rabbit anti-p73 1:1000 (Abcam, ab40658), Rat anti-RFP 1:1000 (ChromoTek, AB_2336064), Rabbit anti-Gad67 1:500 (Nordic BioSite AS, GTX101881), Mouse anti-Calbindin 1:1000 (Swant, AB_3542811), Goat anti-ChAT 1:500 (Merck, AB144P), Mouse anti-TH 1:500 (Millipore, T1299), Goat anti-Serotonin 1:500 (Abcam, ab66047). Next, sections were washed 3 × 10 min in PBS before incubation with secondary antibodies, diluted in incubation solution, for 24h at room temperature (RT). Secondary antibodies: Donkey anti-chicken AF 488 1:500 (Invitrogen, A78948), Donkey anti-Guinea pig AF 405 1:250 (Jackson Immuno Research Lab, 706475148), Donkey anti-Rat AF 594 1:500 (Invitrogen, A21209), Donkey anti-Rabbit AF 647 1:500 (Invitrogen, A31573), Donkey anti-Goat AF 647 1:500 (Invitrogen, A21447), Donkey anti-Mouse AF 647 1:500 (Invitrogen. A32787), Donkey anti-Rabbit AF 568 1:500 (Invitrogen, A10042), Donkey anti-Goat AF 488 1:500 (Invitrogen A32814). After secondary incubation, the sections were washed for a last set of rounds, 3 × 10 min in PBS. The sections were mounted on microscope glass (Thermo Scientific, Superfrost Plus®, 25×75×1) and left to dry on a heating pad. When dry, the sections were washed on the glass with dH_2_O and left to dry again. Finally, the glass was coverslipped (Menzel-Gläser, 24 × 50 mm, #1) with fluoromount (Invitrogen, Fluoromount-G).

For the glycinergic receptor imaging, the protocol is as described above, using sections from RosatdTomatoAi65 +/+; Pde1c-cre +/−; NDNF-flox-FlpO +/− animals (Moan et al., 2025), where CR cells endogenously express tdTomato. Antibodies were as follows: Chicken anti-GFP 1:1000 (Abcam, ab13970), Rat anti-RFP 1:1000 (ChromoTek, AB_2336064), Mouse anti-monoclonal glycine receptor subunit alpha-1 1:500 (Synaptic Systems antibodies, 146 011).

For synaptic puncta staining, the protocol is as described above, using sections from RosatdTomatoAi65 +/+; Pde1c-cre +/−; NDNF-flox-FlpO +/− animals. Antibodies were as follows: Mouse anti gephyrin 1:500 (Synaptic systems, 147011), Rabbit anti PSD95 1:500 (Cell signaling, 2507S).

### Confocal imaging

High-resolution images were acquired with a Zeiss LSM 880 Indimo confocal module attached to a Axio-ImagerZ.2 (Carl Zeiss, Jena, Germany). All images used some common settings throughout all acquisitions. The pinhole size was set to 1 AU, bit depth: 8 bit, and laser intensity and gain was tuned to be within dynamic range. Additionally, pixel dwell was set to shortest duration and averaging at 1. Images were taken with Zen black v.2.1 SP3 (RRID:SCR_013672) and saved in .czi format before being transferred.

### Super-resolution imaging

Super-resolution images were acquired with a Zeiss LSM 990 Airyscan attached to a Axio-ImagerZ.2 (Carl Zeiss, Jena, Germany). All images used the following settings: bit depth: 16, pixel dwell: 0.67 µs, unidirectional scan direction, pixel size: X = 53 nm, Y = 53 nm, Z = 370 nm, acquired using GaAsP-PMT in super-resolution Airyscan mode.

### Scanner imaging

Full series of mounted sections were imaged with a Zeiss AxioscannerZ.1 (Carl Zeiss) controlled with Zen Blue software. Images were acquired with a Plan-Apochromat 20×/0.8 M27 objective, ∼0.33 µm resolution in X and Y, and 4.0 µm in Z. To capture as much as possible from each section, we acquired Z-stacks with 4 slices across 15 µm. Acquired channels: F 405 was excited by LED module 365 nm, with the emission filter set at 412–438 nm; AF 488 was excited by LED module 470 nm, with the emission filter set at 501–538 nm; AF 594 was excited by LED module white with beam splitter 560, with the emission filter set at 570–640; and AF 647 was excited by LED module 625 nm, with the emission filter set at 662–756 nm. These images were saved as .czi files and later split into single channels for further processing.

### Cell counting

The cell counting was done using a modified version of the QUINT workflow (Yates et al., 2019). All sections from one brain were initially imaged using AxioScan scanner. Image files were split into individual sections using the split function in Zen blue. Following, all images were turned into a maximum intensity projection using a script in FIJI/ImageJ (V1.154d, National Institute of Health, USA, https://imagej.net/). The channels of the images were separated to create three groups. One group containing channels with viral signals, namely the channel with GFP signal from helper virus and the channel with RFP signal from RABV. The next group containing only the channel with NeuN signal. Lastly, the remaining group contained only the channel with signal from immunostained p73.

First, the NeuN group was background-adjusted to give the images higher contrast and downsampled such that they all had a width of 1920 pixels. The downsampling was done to fulfill file requirements in subsequent softwares. After the mentioned image processing, the full NeuN group was opened in FastStone Image Viewer v.7.7 (https://www.faststone.org/), which was used to view and put the images in the correct anatomical order. After being organized in the correct order through visual inspection, the files were renamed and numbered to reflect anatomical order. Next, an index file was created for the folder containing all processed NeuN imaged, using the index function in QuickNII (Puchades et al., 2019). The index file prepares the folder to be read into the QuickNII software which was used to align the full series of the brain to the Allen Brain Mouse Atlas CCFv3 (Wang et al., 2020).

The image group with the viral signal from helper virus and RABV was background-adjusted in a similar manner to the NeuN image group, but not downsampled. These were used to count the somas of labeled cells, using the Cell counter plug-in of FIJI/ImageJ (https://imagej.net/plugins/cell-counter). Cells containing only the red fluorophore (input cells) were marked as group one, cells containing only the green fluorophore (non-starter) were marked as group two, while cells containing both fluorophores (starter cells) were marked as group three. All sections were counted and the corresponding coordinates for the counted cells were saved as an .xml file. When all sections were counted, the .xml files were used to create a bitmap mask, in MATLAB (The MathWorks Inc, MATLAB_R2019b), creating a .png image file where each cell coordinate was used to create a pixel of corresponding color to the type of cell. Finally, the MATLAB script saved the .png images in the identical resolution to the downsampled NeuN images. The masks were organized using the FastStone software (FastStone Image Viewer, https://www.faststone.org/FSViewerDetail.htm), in exactly the same order as the NeuN image group.

This concludes all steps needed to quantify cell numbers using the Nutil software, as described in the original publication by Groeneboom and colleagues (2020). Nutil outputs the number of cells, distribution throughout respective brain areas, and the coordinates of each cell. These coordinates were subsequently used to create a visualization of the quantified cells, using Brainrender (Claudi et al., 2021).

Initially, the coordinate file was transformed from .json to .npy, making it readable in a python environment. Following, the array was transformed to the correct mouse voxel coordinates in order for it to fit with the same orientation used in Brainrender. This enabled us to visualize all the counted cells in 3D space over the Allen CCFv3.

### Synaptic puncta quantification

After imaging of synaptic puncta, images were cropped to only contain 1-3 cells and deconvoluted using Huygens deconvolution software (v22.10, Scientific Volume Imaging, The Netherlands, http://svi.nl) with a theoretical point spread function. The background was automatically estimated and double checked for error by the operator. The images were deconvoluted with 100 iterations, a quality threshold of 0.001 and an acuity of 0.0. All puncta in the image were automatically reconstructed in Neurolucida 360 (Micro Bright Field Bioscience), while soma and dendrite were reconstructed with a user-guided method. Next, the resulting data was processed with a python script (v3.11.13; Python Software Foundation, 2023) with an Anaconda environment (v2023.11; Anaconda, Inc., 2023), run in Spyder (6.0.7; Spyder Project Contributors 2023).

All puncta that were within 300 nm from soma and dendrite were kept, while all other puncta were deleted. The resulting puncta number was divided over the soma volume in order to calculate density. Shortly, soma volume was calculated by summing 2D contours from the Z-section images. Lastly, statistical differences between densities were tested using Welch’s t-test. 22 cells at P1 and 26 cells at P15 were evenly sampled from three animals per age group.

### In-vitro electrophysiology

NDNF-flox-FlpO^+/−^;Pde1c-cre^+/−^;Ai65^+/+^ mice of either sex (P4-17) were anesthetized with isoflurane and euthanized with an overdose pentobarbital (IP, 100 mg/kg, Apotekerforeninger). Following, brains were removed and glued to a vibratome-stage, submerged in cutting solution (4 °C) (in mM): 87 NaCl, 2.5 KCl, 1.25 NaH2PO4, 26 NaHCO3, 74 sucrose, 10 glucose, 7 MgCl2, 0.5 CaCl2, saturated with 95% O2 and 5% CO2. In the vibratome (VT1000, Leica), acute hippocampal sections (300 µm) were prepared coronally. Sections were bisected, separating the hemispheres, before being transferred to recover in heated cutting solution (34 °C) for 15 min. After recovery, beaker with section were moved out of the heating bath to cool down to room temperature, where they stayed until recording.

Sections were recorded in a heated (∼35°C) recording chamber of an upright microscope (Zeiss examiner D1, Carl Zeiss, Jena) filled with recording solution, with (in mM): 124 NaCl, 3 KCl, 1.25 NaH2PO4, 26 NaHCO3, 10 glucose, 1.6 MgCl2, 2 CaCl2, saturated with 95% O2 and 5% CO2. Sections were held in place using a platinum ring with nylon strings, and hCR cells were first identified with DIC optics using a 10X (Carl Zeiss, EC Epiplan 10×/0.25 M27) objective. CR cells with a healthy appearance were chosen based on what subfield they resided and ultimately the expression of tdTomato. All cell selections and recordings were performed with a 40x (Carl Zeiss, W N-Achroplan 40×/0.75 M27) objective. Whole-cell patch clamp recordings were done with borosilicate pipettes (Harvard Apparatus, Massachusetts, USA) with a resistance of 6-10 MΩ, filled with an internal solution, with (in mM): 135 CsCl, 0.2 EGTA, 4 Mg-ATP, 0.3 Na-GTP, 10 HEPES, 5 phosphocreatine-Na_2_, 5 QX-314, pH 7.3. 3% Biocytin was added to the internal solution on the same day as recording. All recordings were done with a Multiclamp 700B amplifier (Molecular Devices) controlled by Multiclamp commander 700B, 1550B Digidata (Molecular devices), and Clampex 11 (Molecular Devices). Recordings were sampled at 20 kHz. All cells were held at −60mV or +40 mV when recording.

Protocols and recordings of electrophysiological properties in voltage clamp mode: Excitatory postsynaptic currents events (EPSC, pA) were recorded in gap-free mode for 10-40 min. ACSF with different combinations of modulators (gabazine (12.5 μM), taurine (300 μM, Merck, T0625), strychnine (1.0 μM, Sigma-Aldrich, S8753)) were bath applied into the recording chamber during the recording. Traces were filtered with a low-pass Gaussian 3 kHz filter. Events were identified with template search in Clampfit 11.3 (Molecular Devices), making individual templates per cell. All analysis were either performed with Clampfit 11.3 (Molecular Devices) or with python (v3.11.13; Python Software Foundation, 2023), run in an Anaconda environment (v2023.11; Anaconda, Inc., 2023), with scripts ran in Spyder (6.0.7; Spyder Project Contributors 2023), using PyABF (v2.3.8) to read .abf files from patch clamp recordings. Data was statistically tested using a permutation test for paired samples to investigate if observed amplitude and IEI differences, between ACSF and taurine/strychnine, were greater than chance. Significance levels were set at *: <0.05, **: <0.01.

## Results

### Several hippocampal and extrahippocampal areas target hippocampal Cajal-Retzius cells

To uncover what brain areas and neurons target hippocampal CR cells, we employed transsynaptic rabies tracing. To be able to confidently and specifically target hippocampal Cajal-Retzius cells with the rabies, we took advantage of a CR cell specific transgenic mouse line we previously characterized, the Pde1c-Cre;NDNF-flox-FlpO (fig.1 A; (Moan et al., 2025)). As CR cells express both Pde1c and NDNF, in this transgenic mouse line, they express both recombinases, Cre and FlpO, with the LoxP-flanked FlpO targeted by Cre enabling targeted manipulation of CR cells with FlpO-dependent vectors. We first tested the feasibility of injecting the RABV and helper virus simultaneously in a cocktail injection. P1 Pde1c-Cre;NDNF-flox-FlpO mice pups (fig.1, A) were injected in dorsal hippocampus with the helper:RABV virus mixture (fig.1 B). After perfusion at P15, expression of the respective fluorophores from both RABV and helper virus could be seen confined to hCR cells, showing a clear overlap of expression of both viruses and TRP73 (p73) (fig.1 C-G).

**Figure 1.**
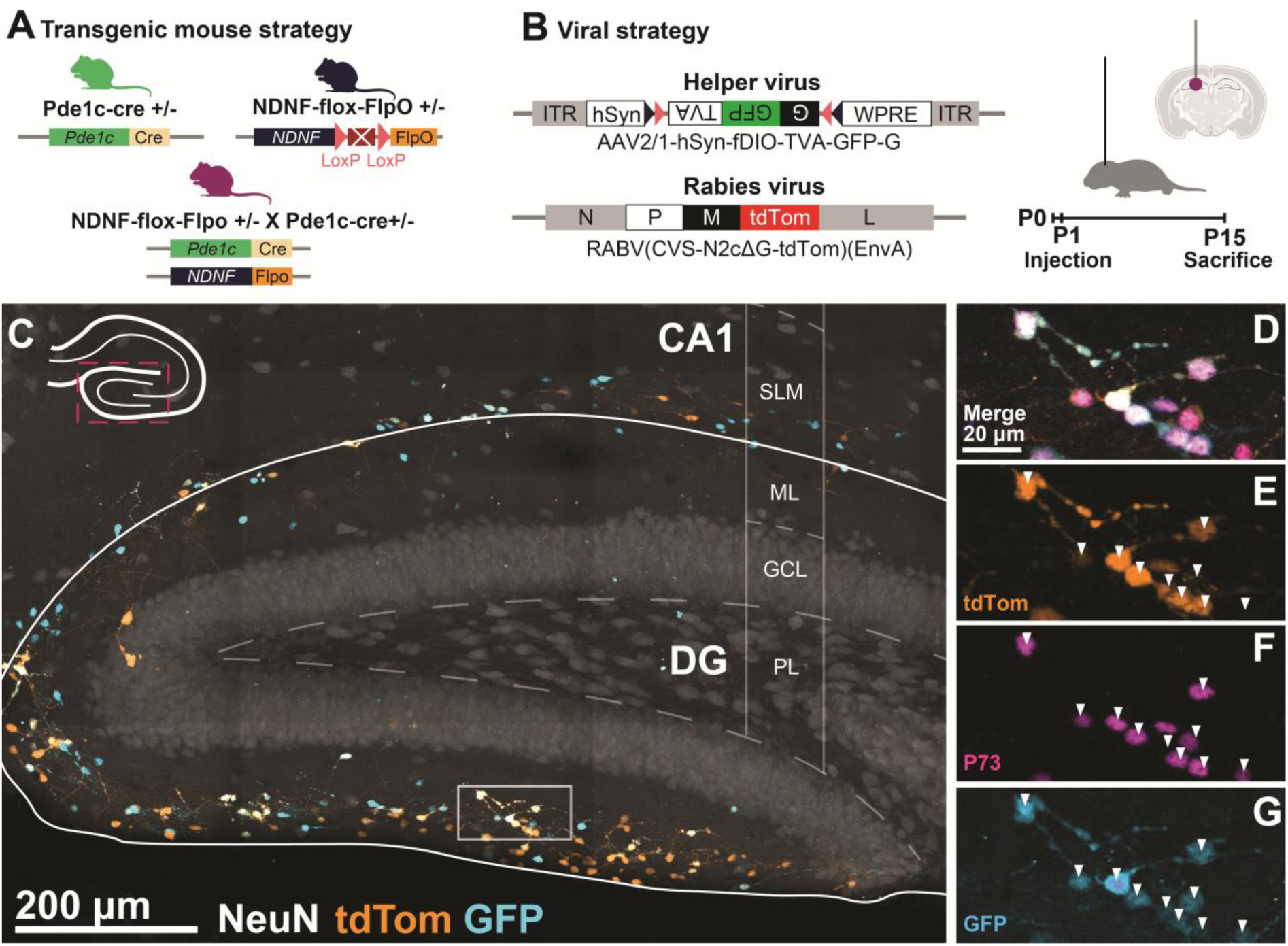
Targeted injection of Cajal-Retzius cells using a Rabies-Helper viral cocktail. **A** Transgenic mouse line created from crossing Pde1c-cre and NDNF-flox-FlpO. **B** Virus injection strategy. Stereotaxic injection was performed at P1 and animals were sacrificed at P15. **C** Image of hCR cells from P15 animal #66502, showing fluorescent signal expressed after transfection of helper virus (orange) and RABV (Cyan). **D-E** Magnified images from inset in **C** showing merge (**D**), tdTomato signal (**E**), p73 signal (**F**), and GFP, arrowheads pointing to cells that have successfully taken up the helper virus (**G**)

With an effective way to infect hCR cells with RABV for retrograde monosynaptic tracing, we started assessing different input populations across the whole brain. Cells only infected with RABV (tdTomato+) and not the helper virus (GFP-) were observed in some cortical, but mostly subcortical areas (fig.2, A). All labeled cells in one full hemisphere were counted in three brains. The labeled cells were segregated into three groups, exclusive GFP+ (helper only), exclusive tdTomato+ (input cells), and GFP+;tdTomato+ (starter cells) (fig.2, A, combination of point clouds from all three brains). The three brains showed a varying number of labeled cells, adding up to *n* = 2510 (#66502), *n* = 14402 (#65217), and *n* = 1902 (#65251) (fig.2, B). The ratio between input cells to starter cells showed a variation between 2.02 for brain #66502, 3.48 for brain #65251, and 2.72 for brain #65217, where the brain with the densest number of labeled cells had a ratio close to the average ratio for all the brains (M = 2.74) (fig.2, C). On average 97.5% of the starter cells were located in the hippocampus, with the remaining 2.5% being located in the neocortex. When checking for overlap between the three labeled cell groups and the CR cell marker p73, we found that on average 88.7% of the starter cells were p73+. The individual percentages for brain #66502 and #65217 were considerably higher, ∼96% and ∼98% respectively. Brain #65251 showed a lower percentage overlap of ∼72%. Furthermore, ∼69% of the non-starter cell group stained positive for p73, indicating that the majority of non-starter cells were also CR cells, and that they had been unable to take up the RABV. Lastly, of all the input cells, ∼30% overlapped with p73 (fig.2, D). In the hippocampus, p73 is almost exclusively expressed in CR cells (Elorriaga et al., 2023; Gumnit et al., 2025; Tissir et al., 2009; Yang et al., 2000) and the input cells overlapping with p73 were only found in hippocampus, suggesting synaptic CR cell-to-CR cell communication in the hippocampus. Despite these variations in densities of labeled cells, ratio of input cells to starter cells, and p73 overlap, the brain regions containing input cells were similar between the three brains.

**Figure 2.**
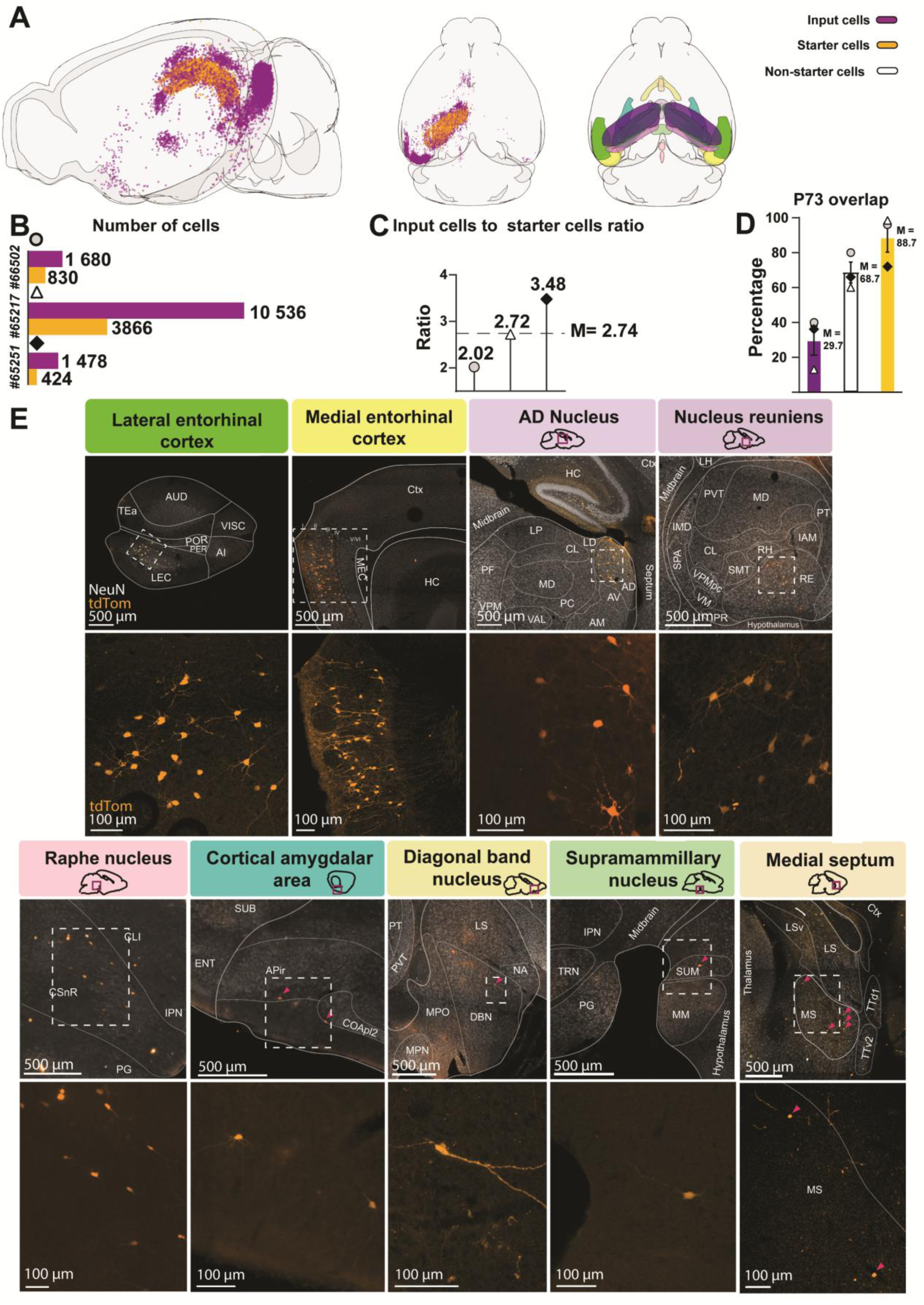
Quantification of virally labeled cells and all observed presynaptic areas. A Left and middle: 3D illustration showing point clouds of starter cells (yellow) and input cells (magenta), combination of all labeled cells from three animals. Right: All presynaptic areas, showed in E. Colored bars indicate categories of cells B Bar chart showing numbers of labeled starter cells and input cells per brain. C Graph showing input cells to starter cells ratio for all brains, average indicated by stippled line, =2.74. D Bar chart for percentage overlap p73 and, from left to right, input cells, non-starter cells, and starter cells. Error bars ± 1 S.E.M, average indicated right of bar, *M.* **E** Images from each extrahippocampal brain area where input cells were located. Top bar showing area name and color matched with 3D illustration from A, and section illustration showing where image was acquired. Top image showing delineated image with input cells (tdtomato+) and NeuN staining. Bottom image, zoomed in area from inset in top image. Scale bars are placed in bottom left of each image.

The input cell populations were distributed across the same regions in all three brains. Populations of different cell numbers were observed in both entorhinal subareas, thalamus, cortical amygdalar area, raphe nucleus, supramammillary nucleus, diagonal band nucleus, and medial septum (fig.2, E). We also collected samples from three injected control animals (double-negative littermates) which showed no expression indicating rabies infection, except for one animal. In this control, a sparse number of subicular pyramidal cells, dentate gyrus granule cells, and cells in the anterodorsal nucleus of the thalamus (ADN) exhibited tdTomato expression (Sfig.1, A-B). Thus, we are not considering ADN as input area onto hCR cells, given the uncertainty of the results. Nevertheless, the number of cells that took up the virus in this one control animal is far fewer than in the experimental animal, and these populations could possibly still be sending inputs to hCR cells. With that said, our experimental paradigm is likely to infect subicular CR cells, which are also potential targets of ADN. This data is still included in the quantification that will follow. No other areas observed in the experimental animals expressed fluorescence in the control.

Populations of input cells could be observed in 7 extrahippocampal areas, in addition to hippocampus CA areas, dentate gyrus, and subicular areas. These results only reflect observed input cells in the ipsilateral hemisphere, but we also observed input cells in the contralateral hemisphere, in LIII MEC and sparsely in the hippocampus (Sfig.2). Of all the extrahippocampal areas with input cells, medial entorhinal cortex (MEC) was the area with the densest population in two (#66502, #65217) out of the three brains (fig. 3, A-C). For the last brain (#65251), dentate gyrus contained the most input cells, followed by thalamus, MEC, and subicular areas, in that respective order. When added together, MEC held the biggest population of input cells, followed by dentate gyrus, thalamus, subicular areas, and LEC (fig.3, C).

**Figure 3.**
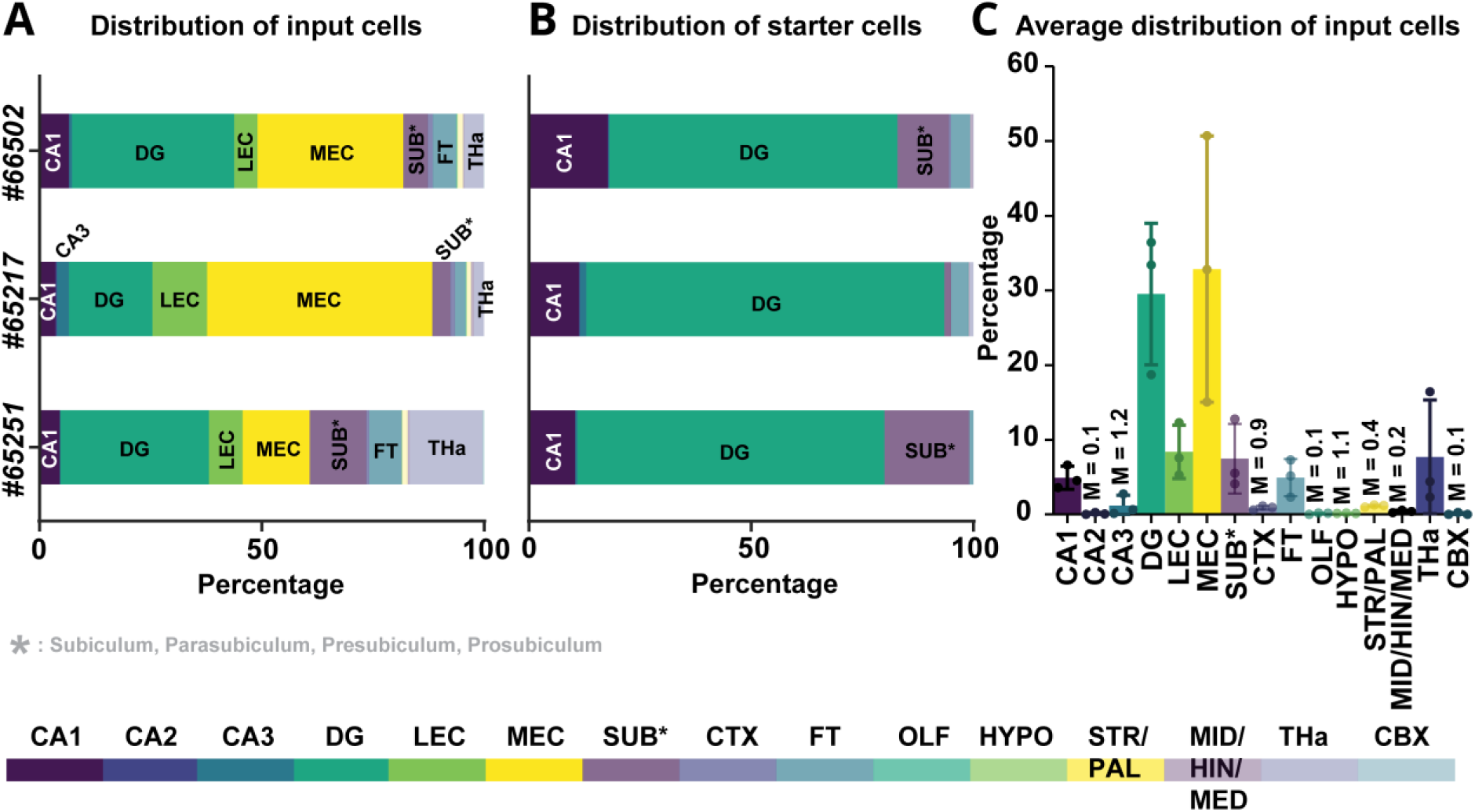
Distribution of input cells and starter cells. A Stacked bar plot showing percentage of input cells in each area, with one bar per brain. B Same as in A, only for starter cells. C Bar plot showing percentage of cells in each area, averaged across all three brains. Individual percentage from each brain indicated by dots, mean indicated by top of bar, M on top of some bars indicate mean value. Overview of all regions included in count, with their respective color.

As EC LII is the major extrahippocampal input to dentate gyrus and CA3, and EC LIII the major extrahippocampal input to CA1(Knierim, 2015; Witter, 2007; Witter et al., 2017), the distribution and density of input we observe onto hCR cells appear to mirror the general pattern of extrahippocampal input specifically into the superficial layers of the hippocampus.

### Hippocampal Cajal-Retzius cells receive predominately putative glutamatergic input

Once we had identified the regions containing presynaptic cells to hCR, we aimed to understand the nature of these cells. Cells of different populations could be observed in the hippocampus and subiculum (fig.4, A). These were sparse numbers of tdTomato+ pyramidal cells in subiculum and in CA1-3, and dentate granular cells (fig.4, A1-4), and, in addition, NGF- and OLM-interneurons (fig.4, B-C). The NGF cells were identified by their location in the SLM and staining positive for GABAergic marker GAD67 (fig.4, B2). The OLM-interneurons were identified by their horizontal morphology and location in CA layer oriens (fig.4, C2).

**Figure 4.**
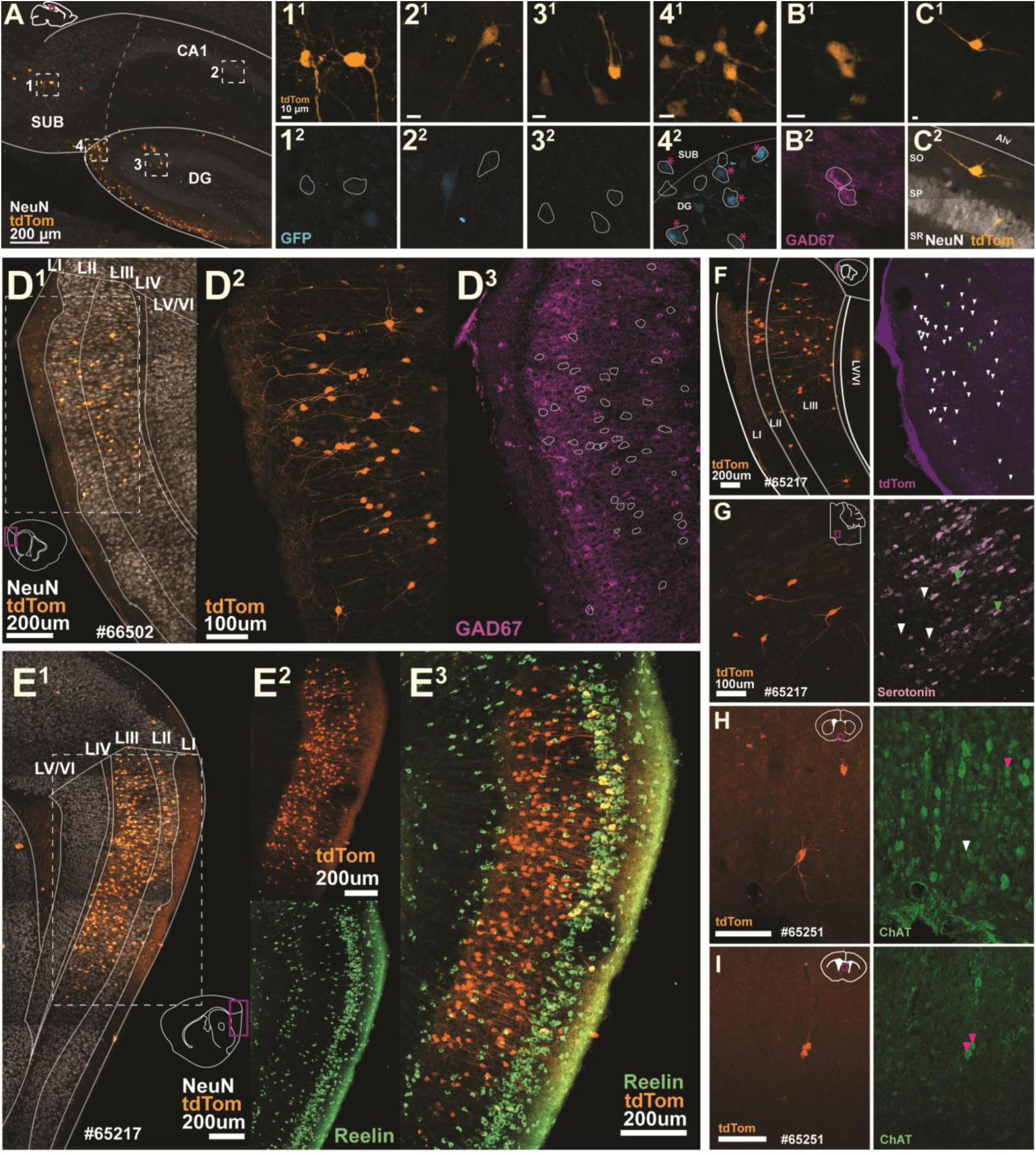
Immunohistological profiles of transsynaptic rabies labeled populations in hippocampus, MEC, raphe nucleus, diagonal band nucleus, and medial septum. **A** Sagittal section from hippocampus showing tdTomato+ cell populations in subiculum (SUB), Cornu Ammonis 1 (CA1), and Dentate Gyrus (DG). **A1-4** Zoomed in areas from insets 1-4 in A. **1^1-2^** Tdtomato+ cells in subiculum and corresponding image showing expression of GFP, illustrating if tdtomato+ cells co-express GFP or not. **2-4** Similar as in 1^1-2^, for CA1, DG, and a selection of CR cells, respectively. Co-expression of tdtomato and GFP indicated with pink asterisk. **B^1-2^** Example tdtomato+ cells imaged from a different hippocampal section, located in SLM, co-expressing GAD67, demonstrating that they are most likely neurogliaform interneurons. **C^1-2^** Putative OLM-interneuron, based on morphology and placement in stratum oriens expressing tdtomato. **D^1^** Sagittal section from animal #66502 showing tdTomato+ cell population in MEC. Section figure bottom left. **D^2^** Magnified area from inset in D^1^ showing tdtomato+ cells. **D^3^** Same image as in D^2^ with GAD67 staining, with outlines of tdtomato+ somas. No overlap between tdTomato+ somas and GAD67 could be identified. **E^1^** Sagittal section from animal #65217, similar to description in **D^1^** with section figure in bottom right. **E^2^** Same section as in E^1^ showing all tdtomato+ cells (top), and reelin staining (bottom). **E^3^** Merge of images in E^2^. All tdtomato+ cells in LII co-expressed tdTomato and reelin. **F** Sagittal section from animal #65217. Left, tdTomato+ cells in MEC. Right, same section with staining for Calbindin. Arrowhead point at all tdTomato+ cells, with green arrowhead indicating overlap between tdTomato and Calbindin. Section figure, bottom left. **G** Sagittal section from animal #65217. Left, tdTomato+ cells in raphe nucleus. Right, same section with staining for serotonin. Arrowheads pointing towards tdTomato+ cells, with green arrowheads indicating overlap between tdTomato and serotonin. Section figure in top middle. **H** Coronal section from animal #65251. Left, tdTomato+ cells in diagonal band nucleus. Right, same section with staining for Choline acetyltransferase (ChAT). Arrowheads pointing towards tdTomato+ cells, pink arrowheads indicating overlap between tdTomato and ChAT. Section inset top middle. **I** same as in H only for medial septum.

In the MEC, all input cells were positioned exclusively in LII-III (fig.4, D). To further investigate the nature of these retrogradely labeled cells we performed immunohistochemical staining for markers used to identify distinctive entorhinal neurons. None of the labeled cells stained positive for GAD67, a marker for putative interneurons, indicating a high likelihood that all cells were principal cells (fig.4, D3). In fact, no cells in the remaining six extrahippocampal areas stained positive for GAD67 (not shown), suggesting that all extrahippocampal input to hCR cells originate from putative glutamatergic cells.

Furthermore, almost all tdTomato+ LII neurons in ipsilateral MEC expressed reelin (Fig.4, E), a marker previously shown to define a specific subset of MEC LII principal neurons, the stellate cell (Varga et al., 2010). In the contralateral hemisphere, no input cells could be observed in LII MEC. In agreement with previous literature describing afferent projections from entorhinal cortex to hippocampus (Steward & Scoville, 1976), all contralateral cells were located in LIII, and these were also GAD67-negative (Sfig.2, A; GAD67 staining not shown). No ipsilateral MEC LII input cells stained positive for calbindin, commonly used to define MEC LII pyramidal cells (Varga et al., 2010), while in ipsilateral LIII a subset of tdTomato+ cells expressed calbindin (fig.4, F).

Outside of entorhinal cortex we examined more closely the input cells in raphe nucleus, diagonal band nucleus, and medial septum. The former area is known to contain serotonergic neurons (Dahlstrom & Fuxe, 1964), and the two latter to contain cholinergic neurons. We therefore decided to stain for these markers to understand the nature of these specific input cells. Indeed, a subset of the tdTomato+ cells in raphe nucleus stained positive for serotonin (Fig.4, G), while a subset of tdTomato+ cells in the diagonal band nucleus and medial septum (Fig.4, H-I), stained positive for choline acetyltransferase (ChAT), a marker commonly used to identify cholinergic neurons (Mesulam et al., 1983). These results demonstrate that hCRs receive neuromodulatory input from the basal forebrain and brainstem.

### Postsynaptic markers decrease with age in hippocampal Cajal-Retzius cells

To reconcile the results from our rabies tracing, showing multiple sources of putative glutamatergic input, we sought to investigate if hCR cells express the proper synaptic anchoring proteins to receive glutamatergic and GABAergic input, and if these persist with age. We therefore examined the expression of postsynaptic markers over hCR cells somas, at two different developmental timepoints, P1 and P15. This would allow us to check for the actual presence of postsynaptic proteins associated with glutamatergic receptors (PSD95) and GABAergic receptors (gephyrin), and how the expression of both PSD95 and gephyrin would change across early postnatal development.

At P1 we observed a dense plexus of both PSD95 and gephyrin around the soma of the CR cells (*n cells* = 22; *n PSD95 puncta = 127.3 ± 13.7; n gephyrin puncta = 116.4 ± 10.2*) (fig.5, A). Both markers were also present at P15, but in noticeably sparser numbers (*n cells* = 26; *n PSD95 puncta = 12.2 ± 2.9; n gephyrin puncta = 35.2 ± 4.7*) (fig.5, B). At the early timepoint, both markers were expressed with a similar density, with no significant difference (*Gephyrin: M = 0.275, SD = 0.095, n = 22*; *PSD95: M = 0.291, SD = 0.127, n = 22; p = 0.6371*). In comparison, at the later timepoint, there was a higher density of gephyrin puncta than PSD95 puncta across the CR cell soma (fig.5, C) (*Gephyrin: M = 0.103, SD = 0.061, n = 26*; *PSD95: M = 0.037, SD = 0.038, n = 26; p <0.0001*). Actually, both the density of gephyrin- and PSD95 puncta had significantly decreased between P1 and P15, but PSD95 more so than gephyrin (*Gephyrin P1 vs P15: p <0.0001*; *PSD95 P1 vs P15: p<0.0001*). This is apparent when looking at the per cell differences in ratio between the two postsynaptic puncta, showing that there was a substantially higher gephyrin puncta density per PSD95 puncta density at P15 compared to P1 (*Ratio P1*: *M = 1.236, SD = 1.196, n = 22; ratio P15: M = 6.478, SD = 6.509, n = 26; p <0.001*) (fig.5, C). This indicates an initially balanced expression of GABAergic and glutamatergic receptors shortly after birth, which eventually is biased towards GABAergic receptors. This could indicate that CR cells primarily utilize glutamatergic receptors prenatally, and GABAergic ones postnatally. Alternatively, there could be a survival bias in CR cells that express less glutamatergic receptors.

**Figure 5.**
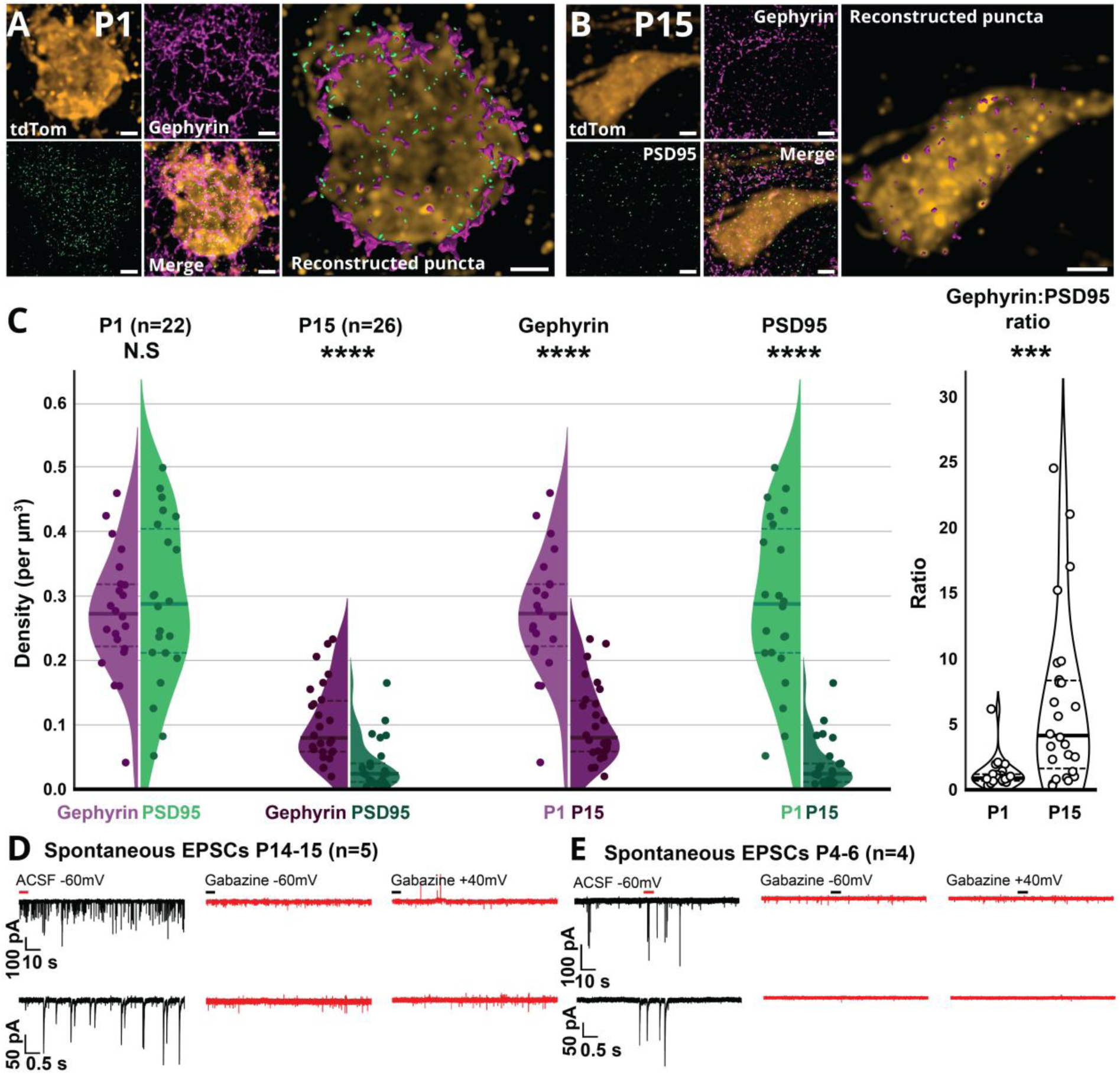
Gabazine fully blocks spontaneous excitatory postsynaptic currents, while CR cells express both PSD95 and gephyrin at both P1 and P15. **A** Super-resolution image of P1 hCR from Pde1c-cre^+/.^;NDNF-flox-FlpO^+/−^;Ai65^+/+^ animals, imaged with four channels showing tdTomato (CR cell, orange) top left, gephyrin (magenta) top middle, PSD95 (green) bottom left, merge bottom middle, tdTomato with reconstructed gephyrin and PSD95 puncta (≤ 300 nm from soma) right. **B** Same as in A, only for hCR from P15 animal. **C** Split violin plots showing density (total number over soma volume) of postsynaptic markers, from left to right, gephyrin and PSD95 at P1, gephyrin and PSD95 and P15, P1 vs. P15 gephyrin, P1 vs. P15 PSD95. Last violin plot shows ratio of gephyrin to PSD95 density at P1 and P15. Significance tested with Welch’s T-test, *** = *P* < 0.001, **** = *P* < 0.0001. **D** From left to right, spontaneous excitatory postsynaptic currents (EPSCs) recorded in CR cells from adolescent animal (P14-15). Top trace, example of CR cell currents recorded in with ACSF at −60 mV, bottom trace, zoomed in portion, indicated by red line in top trace. Middle and left, same as left, only with added 12.5 μM gabazine (red). Recorded at −60 mV (middle) and +40 mV (right). Bottom trace, zoomed in portion, indicated by black line in top trace **E** same as in A, only for young animals (P4-6).

### Gabazine treatment blocks all spontaneous synaptic activity

The surprising amount of observed extrahippocampal glutamatergic input cells led us to reinvestigate the nature of spontaneous synaptic activity in hCR cells. To record glutamatergic synaptic activity, we performed in-vitro patch clamp experiments, recording from hCR cells at −60 mV, as previously done by Quattrocolo & Maccaferri (2013, 2014), with a reversal potential of +1.1 mV for chloride. At P14-15, all spontaneous activity was fully blocked in the presence of 12.5 μM gabazine when clamped at −60 mV. (Fig.5, D). This is corroborated by several other studies, and it is commonly accepted that mouse CR cells show no glutamate receptor mediated spontaneous currents, and minor responses to attempts of glutamate receptor activation (Elorriaga et al., 2023; Marchionni et al., 2010). It has been suggested that these cells might only express certain types of NMDA receptors, and not AMPA (Radnikow et al., 2002), leading to an inability of the NMDA receptor to eject its magnesium ion (Mg^2+^). We then tested the same gabazine treatment while clamping the cells at +40 mV, a membrane potential at which all NMDA receptors would not be blocked by Mg^2+^. Yet, no spontaneous current events were observed in this condition either (Fig.5, D). With our RABV injections happening already at P1, we speculated that the putative glutamatergic synapses were present only at earlier stages of development. We therefore ran the same experiments in tissue from P4-6 mice but observed no spontaneous current events in the presence of gabazine, neither at −60 mV, nor +40 mV (Fig.5, E). Our electrophysiological experiments confirmed previous literature showing an absence of glutamatergic inputs onto CR cells.

### Glycinergic receptors might mediate extrahippocampal input onto CR cells

Previous studies have investigated the possibility of taurine acting on CR cells, given the abundant amount present in the brain at early ages and its role in development (Kilb & Fukuda, 2017; Kilb et al., 2002). PSD95 anchors glutamatergic receptors (gluR) NMDA and AMPA, whose function in CR cells is debated and poorly understood, as mentioned earlier. Gephyrin anchors what we consider inhibitory receptors, GABA_A_, but also glycinergic receptors (glyR) (Kirsch et al., 1993). GABAergic fibers that have been found to contact hCRs, namely NGF and OLM interneurons (Quattrocolo & Maccaferri, 2013), are likely related to the gephyrin expression in hCR cells. Conversely, we assume that the putative glutamatergic input revealed with our rabies tracing likely relates to their expression of PSD95. Still, with the apparent lack of gluR mediated spontaneous EPSCs in CR cells, we sought to consider other possible mechanisms in which communication could be conveyed from glutamatergic fibers. In the hippocampus, a non-canonical synaptic motif of glutamatergic fibers contacting non-glutamatergic receptors, specifically glyRs, have previously been shown (Muller et al., 2013). As such we hypothesized that this could be the motif in question for glutamatergic fibers contacting hCR cells.

To investigate if the input revealed with our rabies tracing could be mediated by activating glyRs instead of gluRs, we labeled hCR cells and MEC LII stellate cells in a 6-week-old Pde1c-Cre^+/−^;Odz3-TTa^+/−^;tetO-GCamp6^+/−^ animal. By using this mouse line, we expressed mCherry in hCR cells with a Cre-dependent vector, and GFP in MEC LII reelin-positive cells with a TTa-dependent GFP. Because of the GCcamp6 expression in this mouse line, all reelin-positive L2 cells in EC will express GFP, additionally, we ensured increased and homogeneous GFP expression in MEC LII reelin-positive cells through an injection with the TTa-dependent vector. This approach allowed us to investigate if fibers from MEC LII stellate cells came into close contact with hCR cells and stained for glyRs to see if these structures are positioned in a way that would facilitate communication. Here, both soma and dendrites of hCR cells express glyRs and came into close contact with the fibers of interest (fig.6, A-B). When reconstructing glyR puncta, we were able to determine that both hCR soma and dendrites have puncta that are positioned within ≤ 300 nm from the membrane (fig.6, B). These puncta again, lay in close proximity (≤ 300 nm) to MEC LII stellate axons (fig.6, C), demonstrating appositions between these three structures, implying probable communication (Hohensee et al., 2008; Schätzle et al., 2012). This was especially interesting for us, indicating that glycinergic receptors, the main target of glycine and taurine, might mediate at least a portion of the extrahippocampal input, and implies that these fibers could make synapses containing glycinergic receptors on hCR cells, instead of presumed glutamatergic receptors. Activation would trigger opening of chloride channels, depolarizing hCR cells in a fashion similar to the response of GABAergic receptors (GABA_A_R) activation (Curtis et al., 1968; Hamill et al., 1983). Indeed, glycine has been shown to be preferentially stored in glutamatergic presynapses, likely co-packaged with glutamate, in the hippocampus. Upon activation of these terminals, they cause strychnine sensitive responses in the postsynaptic site (Muller et al., 2013). Based on the EC LII reelin-positive axon apposition to glyRs in hCR cells, and minimal response to glutamate in CR cells, we argue that non-glutamatergic signaling could be the synaptic signature of long-range, extrahippocampal glutamatergic cells onto hippocampal Cajal-Retzius cells.

**Figure 6.**
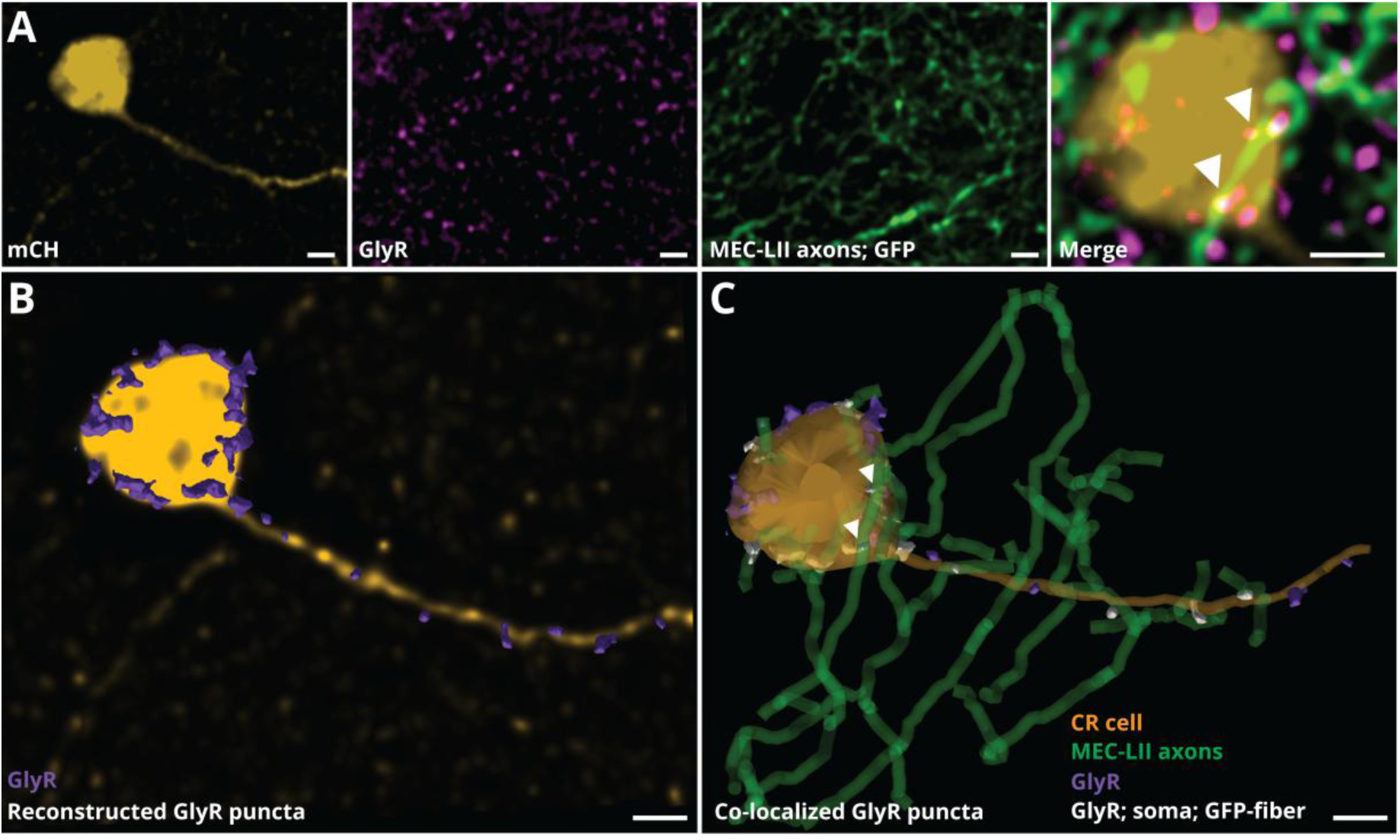
Candidate synapses from EC LII stellate axons position themselves over hCR cell glycinergic receptors. **A** Super-resolution image of hCR and MEC LII stellate axons through cre/TTa-dependent vector injection. From left to right, mCherry expression in hCR (orange), glyR expression (magenta), GFP expression in MEC LII stellate axons (green), merge of all images, white arrowheads pointing at visual overlap creating a white color. Image linearly unmixed to remove potential color-bleed between channels. **B** Reconstructed glyRs that are within 300 nm from CR cells soma and dendrite. **C** Added reconstruction of MEC LII stellate axons that are in proximity to CR glyRs. GlyRs that are both within 300 nm from CR soma or dendrite and MEC LII stellate axon are colored white. Arrowheads point to the two same puncta as in last image of D. All scalebars, bottom right of respective images, 2 µm.

### Glycinergic receptors mediate spontaneous current events in hippocampal CR cells

To further investigate the possibility of glycinergic receptor mediated input onto hCR cells, we performed patch clamp recordings from hCR cells with subsequent bath-perfusion of taurine. Adding 300 µM taurine when clamping hCR cells at −60 mV increased the frequency of current events, which were subsequently fully abolished by adding gabazine (fig.7, A). Taurine caused a slight decrease in amplitude of inward currents that was not significant compared to a bootstrapped shuffle distribution (−18.59%, *p* = 0.2779, *n* = 4) (fig.7, B-C). However, the interevent-interval (IEI) significantly decreased after taurine stimulation, compared to only ACSF (−34.51%, *p* = 0.0047, *n* = 4), indicating an increase in number of events (fig.7, B-C). This demonstrates that taurine significantly increases the occurrence of current events in hCR cells, and that this effect is fully abolished by adding gabazine. To follow up, we wanted to specifically block glyRs using strychnine to test whether hCR cells had spontaneous EPSCs mediated through glyR activation. Bath application of 1.0 µM of strychnine caused the average amplitude of events to drop to almost half compared to events recorded using only ACSF (−42.20%, *p* = 0.0253, *n* = 3). The IEI also increased by 60.65% (*p* = 0.0034, *n* = 3), resulting in fewer events overall (fig.7, E-H).

**Figure 7.**
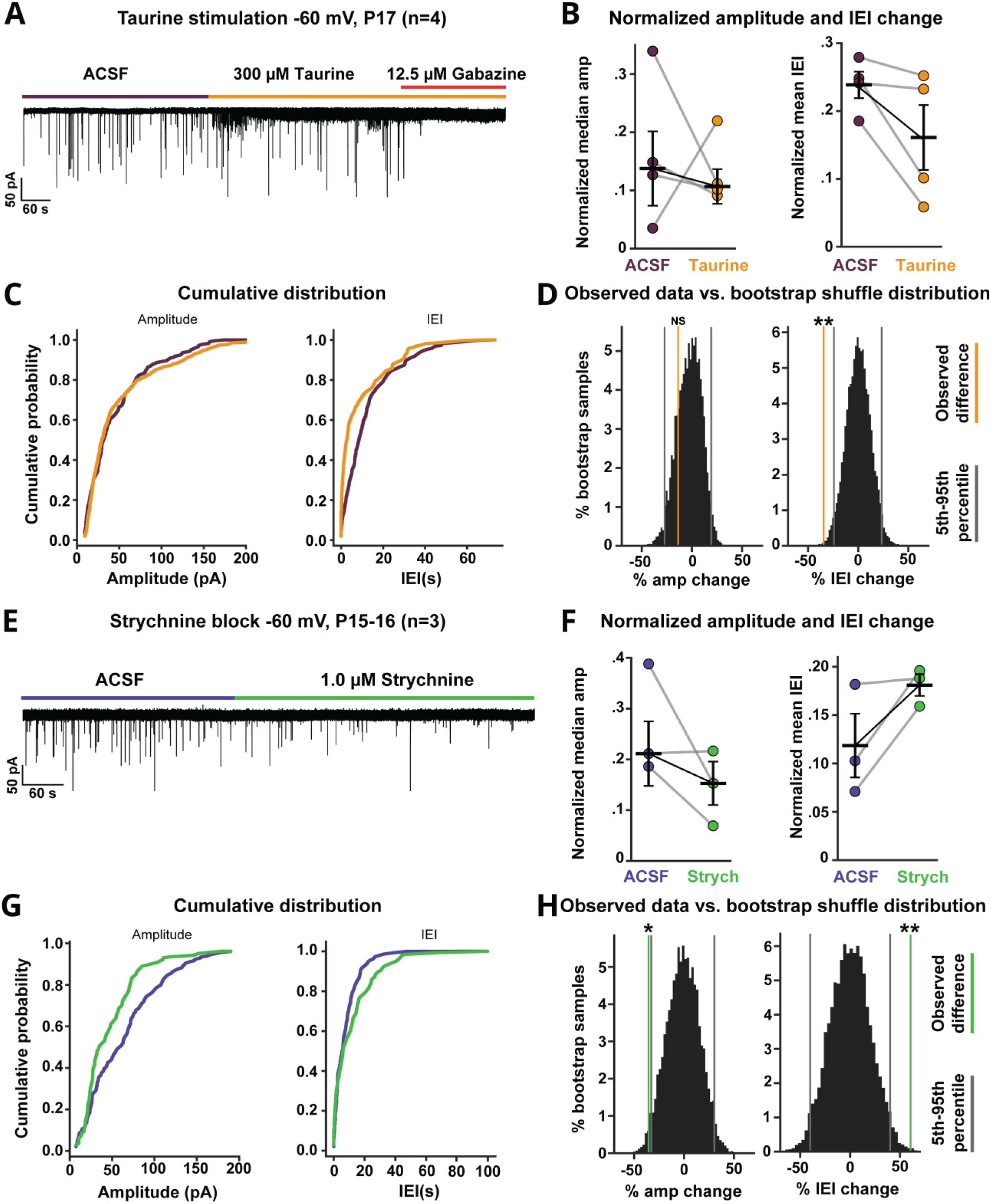
Glycine-receptors mediated spontaneous current events in hCR cells are blocked by gabazine. **A** Example trace from a CR cell from adolescent animal, showing recording of EPSCs after adding 300 μM taurine (orange), and subsequently 12.5 μM gabazine (red). **B** Paired plot of data from individual cells, showing difference in events between ACSF and taurine for amplitude and interevent interval (IEI), respectively. Non-normalized data centered by ACSF grand median (amplitude) or grand mean (IEI). Median/mean represented by bar with respective color. **C** Cumulative distribution plots of non-normalized, pooled data for amplitude (left) and IEI (right) from ACSF (purple) and taurine (orange). **D** Histograms showing distribution of differences between ACSF and taurine, from shuffled, bootstrapped data. ACSF and taurine data were shuffled and bootstrapped individually per cell, before creating an average distribution from all cells, which is illustrated in the histogram. Orange line shows likelihood of the observed difference, compared to average bootstrap distribution. X-axis indicated percentage differences between ACSF and taurine data. **E-H** Same as A-D, only for ACSF vs. strychnine data. *: <0.05, **: <0.01. NS: Not significant

Together, these results point towards the possibility that glyRs mediate CR cell activity, possibly through binding taurine, as shown in these experiments, or glycine.

## Discussion

Cajal-Retzius cells have previously been shown to receive almost exclusively local GABAergic input, however the extent of the connectivity of hCR cells was unknown. To address this, we took advantage of transsynaptic rabies tracing in a CR cell specific mouse line to reveal formerly unknown inputs onto these cells, originating from a large collection of cortical and subcortical structures, as well as locally. Our data shows that (1) hCR cells receive extrahippocampal input from at least 7 different areas; (2) MEC LII stellate - and LIII cells are together the most numerous input population to hCR cells; (3) all input from outside of the hippocampus are exclusively from putative glutamatergic, GAD67-negative cells; (4) GABAergic- and glutamatergic-related postsynaptic puncta are present at hCR cell soma, both decreasing in number with postnatal age, but PSD95 showing a more pronounced reduction; (5) hCR cells express glycinergic receptors that are positioned in physiological plausible distance to incoming MEC LII stellate axons; (6) the number of gabazine-sensitive spontaneous synaptic events increase in hCR cells when stimulated with taurine, and when applying a glycine-receptor antagonist both spontaneous synaptic event amplitude and occurrence decrease.

### Extrahippocampal input onto hippocampal Cajal-Retzius cells

A surprising amount of putative glutamatergic cells, from numerous regions, were revealed through our rabies tracing. This was surprising, as on one hand, spontaneous CR cell synaptic events are sparse and GABAergic input is what drives most, if not all, of this activity (Kilb & Luhmann, 2001; Kirischuk et al., 2014). On the other hand, all areas we observed to send extrahippocampal input onto hCR cells have had their input to hippocampus thoroughly described (Kemppainen et al., 2002; Mesulam et al., 1983; Moore & Halaris, 1975; Van Groen & Wyss, 1995; Vertes, 2015; Witter et al., 2017). The input to hCR cells might be determined by what future cells these inputs ultimately targets. In fact, we did not observe inputs from septal GABAergic cells, which preferentially target other interneurons in both the CA areas and the dentate gyrus, while we detected septal cholinergic cells, that preferentially targets pyramidal cells and granule cells (Amaral et al., 2007; Freund & Antal, 1988). CR cells might mediate the organization of inputs onto principal cells only, which might explain why we only observed cholinergic/GAD67-negative input cells in medial septum.

Several studies have explored the existence of, and responsiveness to, glutamatergic input to CR cells. However, CR cells demonstrate spontaneous synaptic activity only when using high chloride internal solutions, the occurrence of spontaneous synaptic activity does not change in response to gluR blockers (Kilb & Luhmann, 2001), and CR cells show only minor responses to gluR agonist puffing (Marchionni et al., 2010). It quickly appears counterintuitive for at least 7 putative glutamatergic cell populations (fig.2, B; fig.3, E-G) to target a cell type that does not respond to glutamate. Yet, our observations are not the first to indicate that glutamatergic fibers target CR cells. Non-GABAergic synaptic contacts have been demonstrated across cortical CR cells using electron microscopy in juvenile rats (Radnikow et al., 2002), and mice (Anstötz et al., 2014), and local electrical stimulation elicited small AP-5 sensitive responses in CR cells (Kim et al., 1995; Radnikow et al., 2002; Schwartz et al., 1998). Cocas and colleagues (2016) showed with rabies tracing that cortical CR cells receive input from Ctip2+, layer 5 pyramidal neurons. Other studies have described serotonergic (Janušonis et al., 2004; Schwartz et al., 1998), but to our knowledge, not cholinergic input (Schwartz et al., 1998), like we observed from medial septum and diagonal band nucleus (fig.4, H-I).

It is still uncertain whether the inputs we observed onto hCR cells persist following postnatal development, or if they are eliminated when the fibers ultimately synapse onto hippocampal pyramidal cells or dentate gyrus granule cells. We attempted the same tracing experiments at several later timepoints (P15, P21, P30, *data not showed*) but were never able to infect CR cells at any later age. Thus, we cannot confidently demonstrate the existence of these connections at other timepoints. Our results from postsynaptic puncta staining show a significant decrease in both gephyrin and PSD95 between P1 and P15 (fig.5, A-C), indicating that a portion of these connections are eliminated during the first two weeks of development. Indeed, the density of PSD95 decreased more than gephyrin, suggesting that hCR cells go from incorporating a wide range of extrahippocampal input early, to later relying more on local GABAergic, and possibly long-range glycinergic input. Nevertheless, our results also show potential connections between hCR cells and MEC LII axons at 4-6 weeks of age (fig.6), implying the persistence of extrahippocampal inputs even after 1 month. In terms of putative GABAergic contact points, cortical CR cells demonstrate an age-dependent decrease of dendritic vGAT (De Rosa et al., 2026), similar to what we observed for hCR cell somatic gephyrin. This is suggested to result from a local inhibitory circuit transition, wherein CR cell death, and change of target from CR cells to principal cells, are required for proper circuit establishment (Damilou et al., 2024; De Rosa et al., 2026; Riva et al., 2019; Riva et al., 2023)

As an alternative explanation to how glutamatergic fibers could communicate with hCR cells, we hypothesize a synapse of glycinergic nature. We found that hCR cells from animals in the third postnatal week exhibited a decrease in the interevent-interval when stimulated with 300 μM of taurine, triggering more small-to-medium sized current events compared to baseline (fig.7, A-D). Taurine has been observed to activate glyRs at low concentrations (<1 mM) in several brain areas. The exception being the ventrobasal thalamus, where it acts on both glyRs and GABA_A_R at low concentrations (≤ 100 μM), most likely an effect caused by subunit specific interaction of the gamma-less GABA_A_R (Jia et al., 2008). Cortical CR cells are shown to express gamma-subunit GABA_A_Rs after P5 (Cheng et al., 2006). In hippocampal CA1 cells, <1 mM of taurine activated glyRs, while ≥3 mM activated both glyRs and GABA_A_Rs (Wu et al., 2005). Therefore, using 300 μM in our experiments likely avoids GABA_A_R activation in hCR cells. Interestingly, we showed that the effect of taurine in hCR cells is abolished when adding gabazine, a GABA_A_R antagonist (fig.7, A).

As the effect of gabazine and taurine creates a complicated response on hCR cells, we investigated the strychnine sensitivity of spontaneous synaptic activity in hCR cells. 1.0 μM strychnine significantly decreased the amplitude and number of spontaneous synaptic events in hCR cells (fig.7, E-H). This indicates that a considerable portion of the spontaneous activity is driven by activation of glycinergic receptors. Because of this, and the fact that glycine is co-packaged in glutamatergic vesicles in the hippocampus (Muller et al., 2013), we propose that the putative glutamatergic fibers we observed can communicate to hCR cells through glycinergic receptors. This would excite hCR cells and likely cause a feedforward release of glutamate onto SLM NGF-interneurons, as well as ambient release of glutamate into the SLM.

### The role of Cajal-Retzius cells in the hippocampus and their local connectivity

We show that hCR cells receive input from principal cells in subiculum, CA and DG (fig.4, A; Sfig.2, B). This input likely corresponds to CR cells located in the respective subfields of the hippocampus. Furthermore, we confirm findings from Quattrocolo and Maccaferri (2013) demonstrating that NGF-interneurons and OLM-interneurons indeed target hCR cells (fig.4, B-C; Sfig.2, C). Interestingly, we also note CR cell-to-CR cell connectivity, a phenomenon that has been speculated before, but never shown. An input-output relations study of cortical CR cells reported that CR cell axons span remarkably long ranges horizontally across layer 1. These axons preferentially made synaptic contact on non-GABAergic structures, presumably other CR cells and pyramidal cells (Anstötz et al., 2014). We find that hCR cells are synaptically connected to each other, as ∼30% of the input cells in the hippocampus were p73 positive (fig.2, D). This number is probably an underestimation, as most of the hCR cells were starter cells, which means that starter cell CR cells could be passing rabies between each other with no way for us to demonstrate it. Likely, the majority of hCR cells are synaptically connected. Given the glutamatergic nature of hCR cells, the synaptic signature of CR cell-to-CR cell communication could also be mediated by glycine through glycinergic receptors (Muller et al., 2013; Quattrocolo & Maccaferri, 2014).

With the convergency of so many different inputs to hCR cells, and the possible CR cell-to-CR cell connectivity, one could hypothesize how synchronous CR cell activity drives hippocampal development. The extrahippocampal input could activate a subset of CR cells, causing a propagation of depolarization in a network of interconnected CR cells. Recurrent excitation within neighboring neurons as an amplifier has been suggested before, for example for amplification of thalamic input arriving in cortical layer 4 (Douglas et al., 1995). Hippocampal CR cells might be able to amplify weak or noisy input in early development, creating a surge of glutamate output onto hippocampal pyramidal cells and dentate gyrus granule cells. If we pair this with the feedforward excitation circuit of CR cells – pyramidal cells – OLM interneurons (Quattrocolo & Maccaferri, 2014), we find a way in which certain loops will get drastically amplified, especially if the same input arrives on the CR cell and the pyramidal cell simultaneously. This presents a way in which hippocampal CR cells could prime hippocampal networks during early development, before an animal starts executing all relevant behavior or receiving all relevant sensory information (Quattrocolo & Maccaferri, 2014). Possibly, this could also hold true for cortical CR cells, especially as their axons are observed to span very far horizontally, crossing over several brain areas (Anstötz et al., 2014). As such, if CR cell-to-CR cell connectivity will be confirmed in the cortex as well, they are nicely situated to amplify or synchronize activity across brain areas during early development.

If extrahippocampal input to hippocampal CR cells happens through glycinergic mediated activity, it would first create a strong depolarization that could amplify itself through CR cell-to-CR cell recurrency. Eventually, the depletion of chloride in the CR cells would shift the reversal potential of glycine and GABA, given that both transmitters open chloride channels. This is a result of the lack of KCC2 transporters in CR cells (Pozas et al., 2008) which entails that the chloride equilibrium in CR cells are fully dependent on the slow mechanism of NKCC1 transporters (Achilles et al., 2007). Essentially, the physiology of CR cells creates a system where excitation is not necessarily shunted by external inhibition, but instead by a threshold of prolonged activation, which ultimately avoids runaway excitation (Kirischuk et al., 2014; Quattrocolo & Maccaferri, 2013), a motif that possibly would not work if glutamatergic receptors were used.

### Limitations

The primary limitations of our study are the small number of non-specific labeling we observed in one control animal, and our failure to replicate the rabies tracing in older animals. We acknowledge that some of the input areas we report (Subicular pyramidal cells, dentate gyrus granule cells, ADN) must be seen in light of our control results. Nevertheless, the number of neurons expressing both the helper and rabies fluorophore were extremely low in the control, as well as in non-CR cells in the experimental animals. A possible explanation for these control results is leakiness caused by the transgenic mouse-line. Indeed, recombination can occur during diploid stages of gametogenesis, causing sporadic expression in transgene-negative offspring (Song & Palmiter, 2018). Notwithstanding, we are confident that the results we report from experimental animals are valid, especially for all areas not reported in the control animals and that only 1 out of 3 control animals showed expression.

We also observed double fluorophore expression in a few p73 negative cells in the experimental animals, but mostly in one of the three experimental animals. Most of these counts are likely human errors while counting, and not actually cells, but artifacts that fluoresced in both the red and the green channel. Yet, the animal with most non-CR cell starter cells also had the highest number of input cells in ADN, indicating that ADN likely needs to be excluded as an input population to hCR cells.

For our quantification of postsynaptic puncta, we compared density of gephyrin and PSD95. These puncta are clusters of proteins that might result from different numbers of proteins per cluster dependent on it being gephyrin or PSD95. The average size of the puncta also varies between the two proteins. Thus, a direct comparison between number of gephyrin-vs. number of PSD95 puncta might be an inaccurate comparison. Nevertheless, it shows an overall change in density between the two proteins, across age.

In the context of demonstrating apposition between hCR cell glyRs and MEC LII axons, we decided on a threshold of 300 nm to infer a physiologically plausible distance. This was grounded in previous descriptions of similar experiments (Hohensee et al., 2008; Schätzle et al., 2012), but it does not confidently prove its existence. However, Hohensee and colleagues showed that when using a threshold of 300 nm, 83 ± 21% of their identified synapses were confirmed as actual synapses in correlative electron microscopy imaging. Given that we identified >10 hCR cell glycinergic receptors withing 300 nm from MEC LII axons, we find our plausible glycinergic communication hypothesis justified.

Our results using taurine to excite hCR cells align with previously reported gabazine antagonism on glycinergic receptors. Gabazine was found to be a potent antagonist for glycinergic receptors, especially when containing alpha-2 and beta subunits (Li & Slaughter, 2007), which are exclusively those expressed by CR cells (Okabe et al., 2004). This raises questions about the use of gabazine, or formerly used bicuculine, which has an even higher affinity for glyRs (Li & Slaughter, 2007; Wang & Slaughter, 2005), to selectively inhibit GABA_A_R activation. These antagonists have been used to suggest a possible exclusive GABAergic drive in CR cells on several occasions (Cosgrove & Maccaferri, 2012; Kilb & Luhmann, 2001; Marchionni et al., 2010; Soda et al., 2003) using concentrations (>10 μM for gabazine; >30 for bicuculine) at which they also act on glyRs (Li & Slaughter, 2007). With our experimental setup, 12.5 μM gabazine blocked the taurine-mediated activity in hCR cells. With this in mind, we suggest that the role of glycinergic receptors in CR cell activity has been masked through the use of unambiguous antagonists. Strychnine also demonstrates antagonism of GABA_A_Rs at certain concentrations. Hippocampal pyramidal cells are reported to have a half-inhibition dose (IC_50_) of 6.7 μM for strychnine on GABA_A_R-mediated currents (Shirasaki et al., 1991), while 30.0 μM strychnine did not significantly affect GABA induced currents in cortical CR cells from P0-3 aged rats (Kilb et al., 2002). As such, we consider 1.0 μM strychnine selective for glycinergic receptors in hCR cells.

The function of the antagonists used in this study are still being discussed and explored to this day, leading to a certain level of uncertainty in their specificity. For example, strychnine has also been reported to have an IC_50_ of 1.5 µM to GABA-activated currents in spinal cord slices (Jonas et al., 1998), which is lower than what has been reported in the hippocampus (6.7 µM (Shirasaki et al., 1991)). Further, earlier CR cell studies have used a wide range of methods to test antagonist specificity, with for example puffing and bath application, which are likely to affect affinities and needs to be taken into account.

## Conclusion

We reveal that hippocampal Cajal-Retzius cells receive a barrage of extrahippocampal input from putative glutamatergic cells. These fibers position themselves in close proximity to glycinergic receptors expressed by CR cells, indicating that non-glutamatergic signaling could be the signature synapse from glutamatergic cells to hippocampal CR cells. With the strychnine-sensitive portion of hCR cell spontaneous synaptic activity shown here, we argue that this input could drive early hippocampal development through amplification and recurrent excitation, preparing the networks for the activity they need to process in a mature brain. With the results presented here, we expand on the network of hCR cells, arguing that they pioneer a majority of hippocampal input. Further, we propose an extension of what role CR cells play in the hippocampus, and how they mediate input into the hippocampus. Our suggestions will need further work to prove the existence of this glycinergic communication, and we propose target activation of incoming fibers while recording postsynaptic activation in hCR cells. Subsequent application of strychnine would demonstrate its glycinergic nature.

## Acknowledgements

We would like to thank B. A. Zaharia for excellent technical assistance; H. M. Møllergård for genotyping; P. J. B. Girão for assistance with image analysis and code; R.R. Nair for viral preparation and advice. R.H.F. Salih for valuable advice regarding statistics and code; K. Dunville for review; and H. Eneqvist, I.L. Glærum, M. J. Nigro, and all the members of the Quattrocolo lab for their insightful comments throughout this project. We further thank the staff in the animal facility at the Kavli Institute for Systems Neuroscience. The work was supported by a Research Council of Norway (RCN) FRIPRO grants to G.Q. (grant number 324305), RCN Centre of Excellence grants (Centre for Algorithms in the Cortex, grant number 332640; Centre of Neural Computation, grant number 223262; to G.Q.), a NARSAD Young Investigator Grant from the Brain and Behavior Research Foundation to G.Q. (grant number 31551), and the Kavli Foundation (to G.Q.). The experiments were performed at the NORBRAIN Facility, Norwegian University of Science and Technology (NTNU) (grant number 295721).

## Authors contribution

Conceptualization: KM and GQ; Data curation: KM; Formal analysis: KM; Funding acquisition: GQ; Investigation: KM; Methodology: KM and GQ; Visualization: KM; Writing - review & editing: KM and GQ.

## Supplementary

**SFigure 1.**
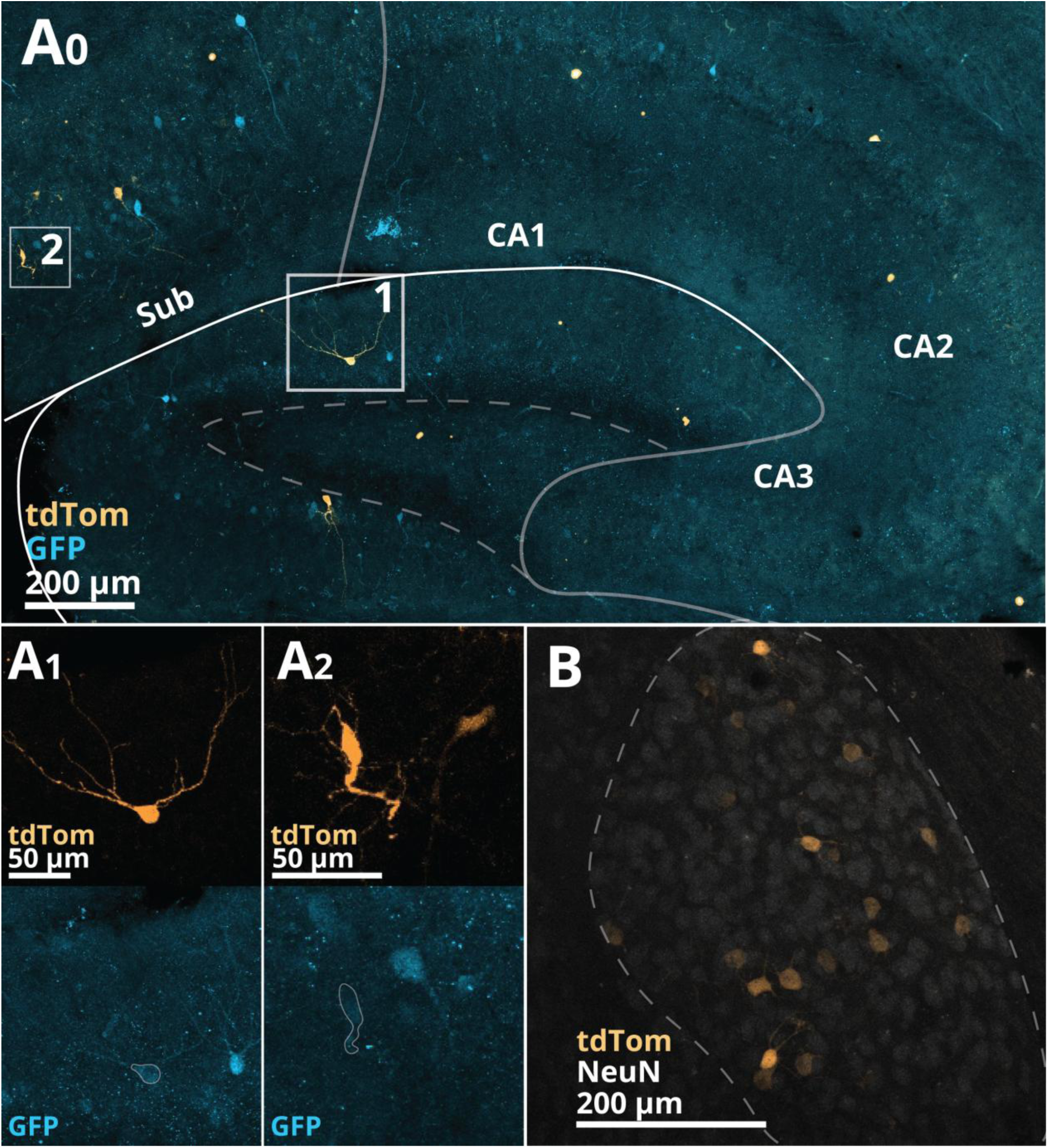
Confocal image from the one control animal showing rabies-expressing cells. **A** Sagittal section of hippocampus showing sparse expression of both tdtomato and GFP in subicular pyramidal cells and dentate gyrus granule cells. **A1** Magnified image of granule cell, showing expression of tdtomato from rabies (top) and the corresponding GFP channel (bottom). **A2** Same as in A1, only for subicular pyramidal cell. **B** Anterodorsal nucleus of thalamus, showing tdtomato+ cells as in experimental animals. These cells were GFP-negative, and the only other population of cells (together with the sparse population in hippocampus from A) that expressed fluorescence in both experimental and control animals. Therefore, we cannot confidently say that ADN, subicular pyramidal cells, or dentate gyrus granule cells are hCR cell input cells.

**SFigure 2.**
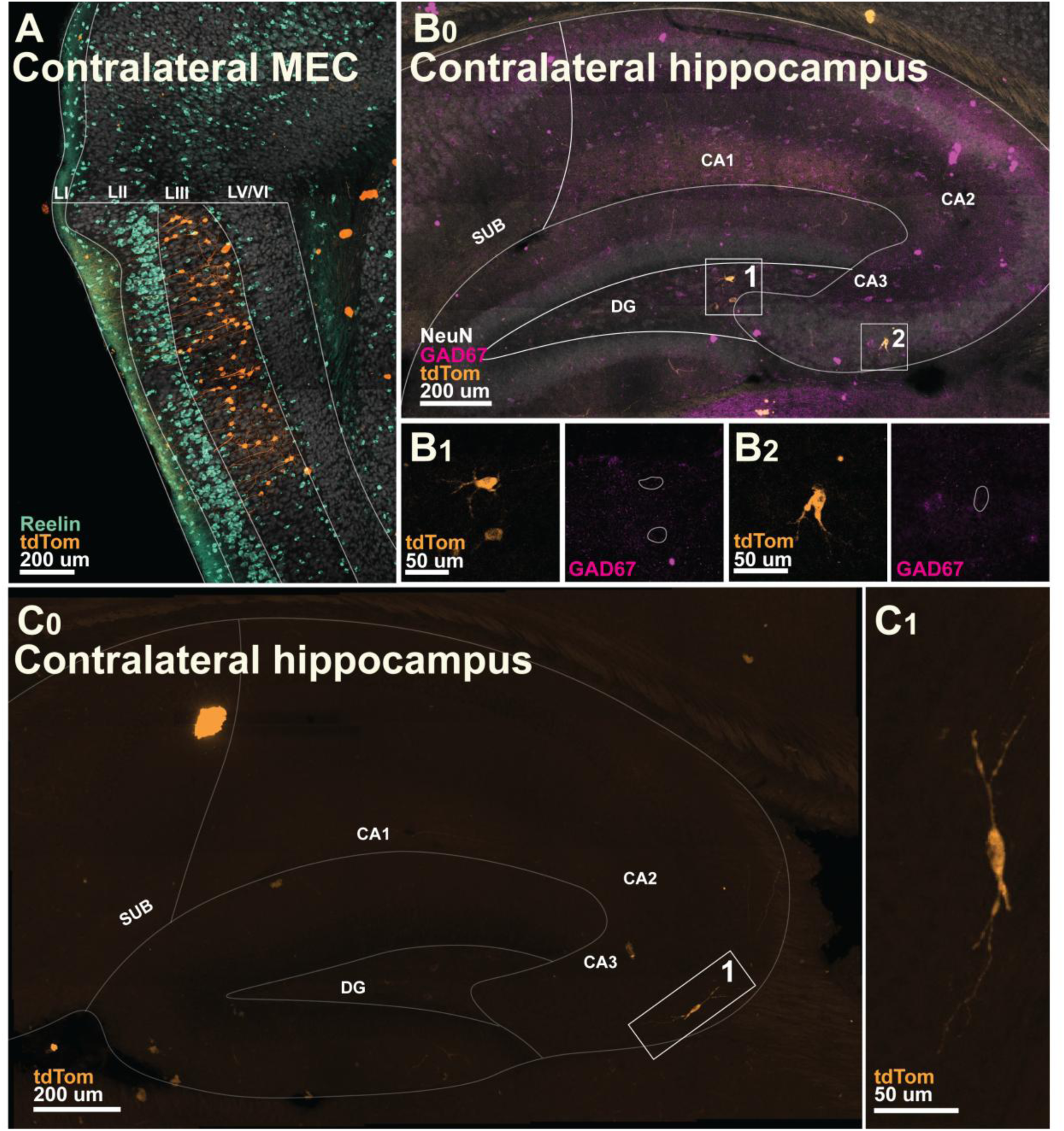
Confocal image from contralateral sections of #65217, experimental animal. **A**. tdTomato expression could only be observed in LIII of contralateral MEC, as opposed to LII and LIII ipsilateral. **B** Sparse number of dentate gyrus mossy cells and CA3 pyramidal cells were observed in the contralateral hippocampus. These were confirmed not to be GABAergic neurons through GAD67 staining. **B1** Magnified image from **B0** showing tdtomato expression in dentate gyrus mossy cells (left) and staining for GAD67 (right). Highlights of soma show that none of the cells expressed GAD67. **B2** same as in B1, only for CA3 pyramidal cell. **C** Sparse number of OLM-interneurons were observed in contralateral hippocampus. **C1** magnified image from inset in C0. Identity of cell confirmed through layer position and stereotypical morphology.

